# MapZ forms a stable ring structure that acts as a nano-track for FtsZ treadmilling in *Streptococcus mutans*

**DOI:** 10.1101/218677

**Authors:** Yongliang Li, Shipeng Shao, Xiao Xu, Xiaodong Su, Yujie Sun, Shicheng Wei

**Affiliations:** Central Laboratory/Department of Oral and Maxillofacial Surgery, School and Hospital of Stomatology, Peking University; National Engineering Laboratory for Digital and Material Technology of Stomatology, Beijing 100081, China; Biodynamic Optical Imaging Center (BIOPIC), School of Life Sciences, Peking University, Beijing 100871, China

**Author notes:** Corresponding Author Correspondence: Shicheng Wei^1^ or Yujie Sun^2^, 1. Central Laboratory/Department of Oral and Maxillofacial Surgery, School and Hospital of Stomatology, Peking University, 22 Zhonguancun South Road, Haidian District, Beijing 100081, China; E-mail address, 2. State Key Laboratory of Biomembrane and Membrane Biotechnology, Biodynamic Optical Imaging Center (BIOPIC), School of Life Sciences, Peking University, 5 Yiheyuan Road Beijing 100871, China.

**Keywords:** Cell division, FtsZ treadmilling, MapZ, Single molecule tracking, Super-resolution imaging, *Streptococcus*

## Abstract

Bacterial binary division requires the accurate placement of the division machinery. FtsZ, the vital component of the division machinery, can assemble into filaments and self-organize into a ring structure (Z-ring) at the proper site for cell division. Thus, understanding how bacteria control the spatiotemporal formation of the FtsZ ring is crucial for small molecule and nanoparticle antibacterial drug discovery. MapZ, a recently identified FtsZ regulator in *Streptococcaceae,* has been found to localize at the mid-cell and position the FtsZ ring. However, the mechanism is still unclear. Here, by using total internal reflection fluorescence microscopy, super-resolution imaging, and single molecule tracking, we investigated the mechanism by which MapZ regulates the FtsZ ring position. The results show that FtsZ exhibites dynamic treadmilling motion in S. *mutans.* Importantly, depletion of MapZ leads to an unconstrained movement of treadmilling FtsZ filaments and a shorter lifetime of the constricting FtsZ ring. Furthermore, by revealing that MapZ forms an immobile ring-like nanostructure at the division site, our study suggests that MapZ forms a stable ring that acts as a nanotrack to guide and restrict treadmilling FtsZ filaments in S. *mutans*, representing a novel way in which bacteria control the division.

---

In bacteria, the eukaryotic tubulin homolog FtsZ^1^ is a core component of the division machinery. On the initiation of cell division, FtsZ self-organizes into a ring (Z-ring), a heterogeneous nanostructure that localizes to the middle of a cell^2^ and recruits other components to construct the division machinery.^3^ Since the extremely vital role of Z-ring for bacterial division, a number of studies have tried to understand the mechanism of FtsZ regulatory system that controls the positioning and timing of Z-ring formation. In well studied model bacterial such as *Escherichia coli* and *Bacillus subtilis,* the Min system^4^ and the Nucleoid Occlusion (NO) system^5^ are two main negative regulatory systems, which prevent the Z-ring assembly at an improper position by inhibiting FtsZ polymerization.^6–7^ MipZ is another negative FtsZ regulatory system, which was discovered in *Caulobacter crescentus* and is conserved among *α proteobacteria.* It directly interacts with FtsZ and alters the conformation of FtsZ filament to prevent improper FtsZ ring formation.^8^ Distinct from the negative regulatory systems mentioned above, several positive control systems were also reported. For instance, SsgA/SsgB- and PomZ-mediated system were identified in *Streptomyces* and *Myxococcus xanthus,* respectively.^9–10^ These systems do not affect the position of Z-ring by inhibiting the improper FtsZ polymerization. Inversely, they localize at the division site prior to FtsZ and regulate FtsZ ring formation at the mid-cell.

Recently, another important positive regulatory system was discovered in *Streptococcus pneumoniae* and is conserved in *Streptococcaceae*, which lack the aforementioned FtsZ regulatory systems. This new system is mediated by the transmembrane protein MapZ (also called LocZ), ^11–12^ which is a substrate of the Ser/Thr kinase StkP (or called PknB^13–14^) and binds to peptidoglycans *via* the extracellular domain.^11, 15^ MapZ positions the Z-ring by directly interacting with FtsZ *via* the cytoplasmic domain. In the absence of MapZ, FtsZ ring mislocalized at the mid-cell and showed an aberrant plane angle.^11, 16^ Notably, differing from other identified FtsZ regulators such as MinC, SimA, MipC and SsgB, which were confirmed to directly interact with FtsZ and exhibit an impact on FtsZ polymerization so as to regulate the position of Z-ring^,6–8, 10^ MapZ does not show remarkable influence on the polymerization or GTPase activity of FtsZ.^11^ This suggests that MapZ regulates the FtsZ position in a novel way, that has not been reported. Despite this important role of MapZ, few studied the mechanism by which MapZ regulates the Z-ring position.

Furthermore, FtsZ can self-organize into filaments, which exhibits a dynamic treadmilling *in vitro*.^17^ Recently, this dynamic treadmilling was experimentally confirmed in model bacteria, *E. coli* and *B. subtilis.^18–19^* The treadmilling of FtsZ filaments drives the cell wall synthesis and is crucial for septum formation and cell division. To gain more insight into how MapZ regulates the FtsZ position, we, therefore, studied the relationship between MapZ and the treadmilling of FtsZ. We show that MapZ is crucial for the proper cell shape and localization of FtsZ ring in *S. muatans*. It controls the localization and plane angle of FtsZ ring by affecting the dynamic treadmilling of FtsZ filaments. Further, by using super-resolution imaging and single molecule tracking, we discover the MapZ forms a stable heterogeneous ring-like nanostructure at the division site, which acts as a track to guide and restrict treadmilling FtsZ filaments to form a proper Z-ring. Our study sheds light on the mechanism by which MapZ control the FtsZ ring position in *Streptococcaceae.* Importantly, it also gains an insight into understanding how bacteria regulate the cell division.

## Results and Discussion

Here, to gain more understanding of how MapZ controls the position of FtsZ ring, we started with a MapZ homolog in *S. mutans*, an important gram-positive pathogenes and easy access to cultivation, genetic manipulation.^20^ We identified the MapZ homolog in *S. mutans* (smMapZ); smMapZ shows only 37% sequence identity with MapZ in S. *pneumoniae* (spMapZ), with 32% and 50% sequence identity for the cytoplasmic and extracellular domains, respectively. Note that, the low sequence identity of spMapZ in *S. mutans* is consistent with previous report,^21^ which showed that average sequence identity of the cytoplasmic domain of spMapZ is 28.4% in *Enterococcaceae* and *Steptococcaceae* while that is 59.5% for the extracellular domain. Most of the first 40 amino acids of spMapZ, which were thought to be sufficient for the interaction of spMapZ and FtsZ, were deficient in smMapZ while the two phosphorylation sites^11–12^ (T67 and T78) and the amino acids^15^ (R409, Y411, N428, Y430, Y450, F451, and N454) required for binding to pneumococcal peptidoglycans are conserved (Figure S1A). Subsequent prediction of sequence disorder tendency (Figure S1B), amino acid conservation (Figure S1C) and analysis of the secondary structure of the intracellular domain (Figure S1D) and suggested that smMapZ forms a similar potential structure to spMapZ, both contain a predicted functional subdomain at the N-terminus of the cytoplasmic domain and two subdomains of the extracellular domain.

smMapZ is essential for regular cell shape formation. Deletion of *smmapZ (ΔsmmapZ)* led to shorter, irregularly shaped cells and mini-cells (Figure 1A, B), which is similar to the phenotype in S. *pneumoniae.* Then, to assess the localization pattern of smMapZ, we fused enhanced green fluorescent protein (EGFP) to the N-terminus of smMapZ under the control of the native promoter. Time-lapse imaging of EGFP-smMapZ (Figure 1C) indicated that smMapZ formed a ring at the mid-cell and a pair of rings appeared as the cell elongated. The distance between the two smMapZ rings increased as a function of cell length while the distance between the smMapZ rings and their nearest cell pole did not change remarkably (Figure 1D), suggesting that the smMapZ ring at the mid-cell split into two rings, which moved to the cell equator of the new daughter cells. Interestingly, smMapZ with cytoplasmic domain deletion *(smmapZΔcyto)* resulted in loss of the ability to position at the constricting septum, resulting in distribution throughout the cell membrane, whereas the extracellular domain null mutation *(smmapZΔextra)* retained a septum localization pattern (Figure 1E). Further multi-domain truncation analysis revealed that the predicted subdomain at the N-terminus of the cytoplasmic domain and the subdomain tandem to the transmembrane domain respond to such localization patterns (Figure S2). This result differs from that of spMapZ.^11^ Try to explain this, we performed an evolutionary tree analysis and revealed that smMapZ and spMapZ belong to different evolutionary branches (Figure S3), which indicated that *S. mutans* might adopt another way in which smMapZ localizes at the septum. The sequence analysis and characterization of smMapZ indicated the similar structure and role in regulating FtsZ ring of smMapZ and spMapZ.

**Figure 1.**
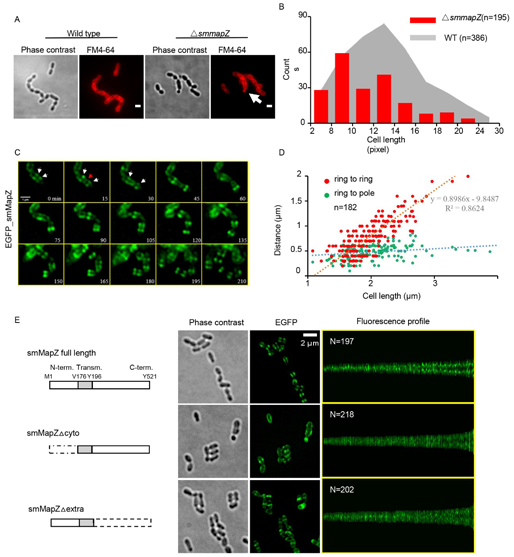
Characterization of a MapZ homolog in *S*. *mutans* (smMapZ). (A) Cell shape of wild type (WT) and deletion of *smmapZ (ΔsmmapZ).* Membrane was stained with FM4-64. A minicell is indicated by the white arrow. Scale bar, 1 μm. (B) Analysis of cell length of WT and *ΔsmmapZ.* (C) Live cell imaging of enhanced green fluorescent protein (EGFP) smMapZ. White arrows indicate the split smMapZ rings, and the red arrow indicates the third smMapZ ring. (D) Analysis of the distance between the two outer rings, and between the ring and the nearest pole. (E) Localization pattern of WT smMapZ, smMapZ with cytoplasmic domain deletion (smmapZΔcyto), and the extracellular domain null mutation (smMapZAextra). A schematic of the mutations is shown on the left, cell shape and EGFP signals in the middle, and corresponding maps of fluorescence intensity (sorted by cell length) on the right.

Indeed, consistent with a previous report,^11–12^ FtsZ and smMapZ were colocalized at the mid-cell in a newborn cell (Figure 2A). Moreover, smMapZ was required for correct localization of the FtsZ ring, as 64% of FtsZ rings were unable to localize at the mid-cell in the *ΔsmmapZ* strain, while only 22.7% of FtsZ rings in the wild-type (WT) strain were mislocalized (Figure 2B). As the cell grew, the smMapZ ring split into a pair, which moved toward the cell equator of the daughter cells before the FtsZ ring. A third smMapZ ring emerged at the division site, constricting with the FtsZ ring. Subsequently, new FtsZ rings assembled, colocalizing with the smMapZ rings of daughter cells.

**Figure 2.**
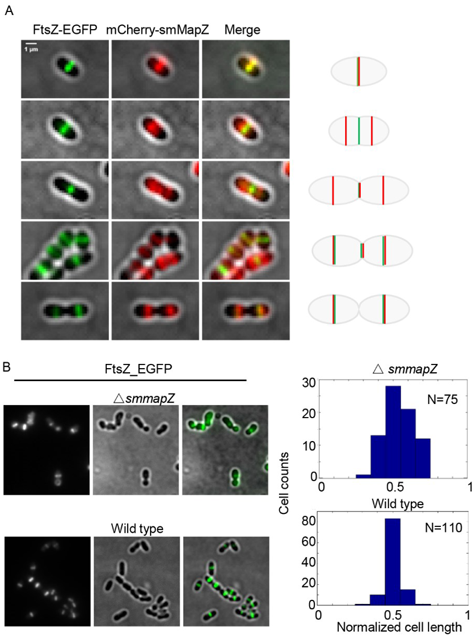
Effect of smMapZ on FtsZ localization. (A) Dual color imaging of smMapZ and FtsZ in different stages of the cell cycle. (B) Localization of FtsZ in WT and *ΔsmmapZ.* Cell shape observed by phase contrast microscopy and fluorescence of EGFP are shown on the left, and statistical analysis of FtsZ ring position is shown on the right.

As a tubulin homolog in prokaryotic, the treadmilling motion of FtsZ filaments was confirmed *in vivo* recently.^18–19^ MapZ did not affect the polymerization and GTPase activity of FtsZ,^11^ but deletion of MapZ did impact FtsZ ring localization and constriction. Thus, we proposed that MapZ might control the treadmilling of FtsZ filaments. To confirm this hypothesis, we labeled FtsZ with a brighter green fluorescent protein, mNeongreen, and used TIRFM to study the dynamics of FtsZ rings. The results revealed a periodic intensity fluctuation of the FtsZ ring (Figure 3A, Figure S4A). With kymography, we observed directional movement of the FtsZ ring (Figure 3B, Movie s1), similar to that in *E. coli* and *B. subtilis,^18–19^* indicating that the treadmilling behavior of FtsZ filament is conserved in different bacterial species. In contrast with *E. coli,* in which only one FtsZ ring forms during a cell cycle, three FtsZ rings (dumbbell structure) could be observed in the majority of *S. mutans* and *S. pneumoniae^11–12^* cells at a later stage of the cell cycle. Surprisingly, we observed translocation of FtsZ filaments from the middle (old) FtsZ ring to the nearby (new) FtsZ rings (Figure 3C, Movie s3) that has not been reported in other bacterial species. Considering that the speed of translocation was similar to that of treadmilling inside the ring and the translocated FtsZ would participate in the new ring formation, it is reasonable to hypothesize that such kind of FtsZ translocation is a functional process rather than abnormal localization. The treadmilling and translocation of FtsZ also support the view that the FtsZ ring consists of multiple protofilaments, thereby demonstrating a bead-like structure or discontinuous clusters.^22–26^

**Figure 3.**
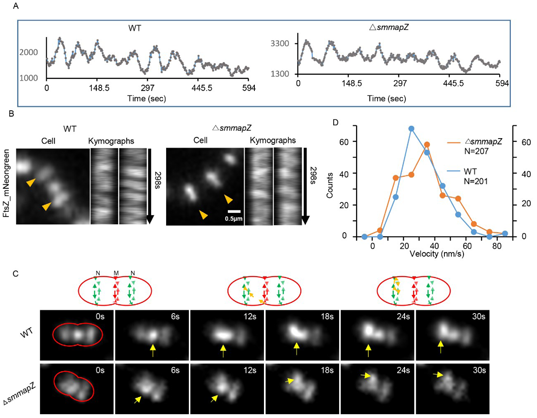
FtsZ shows directional movement independent on smMapZ. (A) Typical periodic fluctuation of the FtsZ ring in WT and *ΔsmmapZ*. (B) Typical direct movement of FtsZ in WT and *ΔsmmapZ*. (C) Distribution of velocity of FtsZ movement inside the rings. (D) Translocation of the FtsZ filament. Schematics are shown above. “M” refers to the middle (old) ring and “N” refers to newly formed rings. Fluorescence images are shown below. The red line indicates cell shape, and yellow arrows mark the initiation of translocation.

Interestingly, deletion of *smmapZ* led to highly unconstrained movement of FtsZ filaments (Figure 4A, Movie s5, s6), but did not perturb the treadmilling and translocation of FtsZ filaments (Figure 3A-C, Movie s2, s4) or alter the speed of treadmilling (Figure 3D), which suggests a crucial role of smMapZ in guiding the treadmilling movement of FtsZ filaments. Although the unconstrained FtsZ filaments could still form a ring-like structure as the cell grew, the angle of the FtsZ ring plane changed back and forth during FtsZ ring formation, which might ultimately lead to aberrant localization of the Z ring. Nevertheless, once the Z-ring formed, the angle of the Z-ring plane would no longer change (Figure S4B, Movie s5). These data can explain why depletion of MapZ leads to an irregular cell shape and minicell formation.

**Figure 4.**
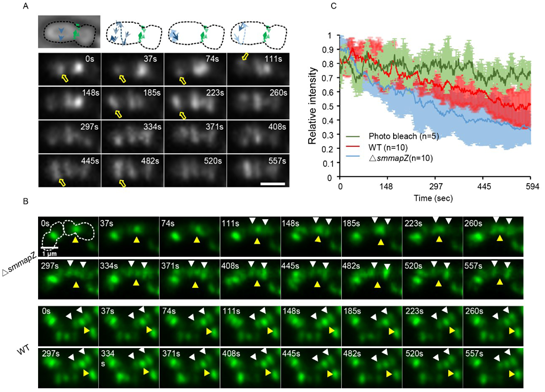
Deletion of *smmapZ* lead to unconstrained FtsZ treadmilling and shorter lifetime of constricting FtsZ ring. (A) Unconstrained movement of the FtsZ filaments in *ΔsmmapZ.* Diagrams of the first four snapshots are shown in the upper panel, and the cell shape in bright field, marked by a dotted line, is shown in the upper left. In the diagrams, the dark blue arrows indicate the position and direction of movement of FtsZ filaments, and light blue arrows indicate the movement path of FtsZ. In snapshots of the time-lapse imaging shown below, yellow arrows mark the position of FtsZ and the change of plane of the FtsZ ring. (B) Snapshots of the time-lapse imaging of three FtsZ ring structures in WT and *ΔsmmapZ*. White arrows mark the newly formed FtsZ ring position and yellow arrows show the position of the middle (old) FtsZ ring. (C) Fluorescence intensity traces of the middle (old) FtsZ ring as a function of time.

Consistent with previous study,^11^ we also noticed that deletion of *smmapZ* resulted in a decrease in chance to observe cells with three FtsZ rings (Figure S4C). By monitoring the complete process of emergence and disappearance of the three FtsZ ring structure, we discovered that the middle FtsZ ring in the *ΔsmmapZ* strain only remained 5-6 min after new FtsZ ring formation, exhibiting a shorter lifetime than the life time (>10 min) in the WT strain (Figure 4B, Movie s7, s8). Analysis of the intensity trace also indicated that deleting *smmapZ* led to a faster fluorescence decrease of the middle FtsZ ring than that in WT (Figure 4C). We speculate that the shorter lifetime was caused by an increase in FtsZ translocation (Movie s5) in absence of the restricting and guiding role of smMapZ; however, we cannot exclude the possibility that it may have been due to premature constriction of the FtsZ ring, as previously described.^11^

Next, to understand how smMapZ guides the treadmilling of FtsZ filaments, we performed three-dimensional structured illumination microscopy (3D-SIM) to study the nanostructure of a Halotag-fused smMapZ (HalojF549-smMapZ^27^) ring. The smMapZ ring exhibited a heterogeneous ring-like nanostructure at the division site (Figure 5A-C, Movie s11), similar to previously studied FtsZ, EzrA, Pbp2^23^, FtsA and ZipA^25^. Such discontinuous ring nanostructures allow for direct observation of their directional movements using single molecule TIRF imaging.^18–19^ Here, to study the dynamics of smMapZ ring, we labeled smMapZ with mNeongreen for TIRF imaging. Interestingly, no remarkable periodic motion of smMapZ was observed by intensity traces or kymograph analysis (Figure 5D, Figure S5A, Movie s9), suggesting that the smMapZ ring is immobile. We further observed the relative movement between smMapZ and FtsZ using a dual color assay. By labeling smMapZ and FtsZ with EGFP and mScarlet-I,^28^ respectively, we revealed that the FtsZ ring exhibited directional movement while the smMapZ ring showed no remarkable movement (Figure 5E).

**Figure 5.**
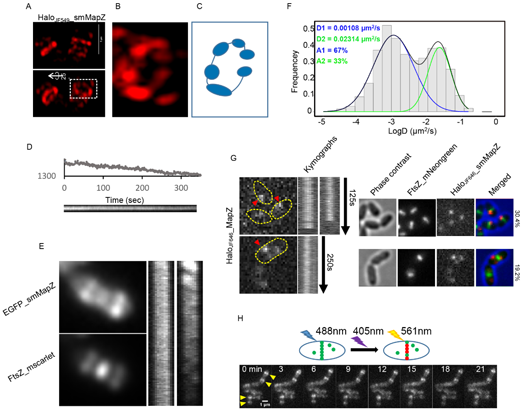
Characterization of the smMapZ ring. (A) Three-dimensional structured illumination microscopy of HaloJF_549__smMapZ. (B) The heterogeneous ring structure of smMapZ in the box region of (A). (C) Cartoon of the heterogeneous ring structure in (B). (D) Intensity trace and kymograph analysis of mNeongreen_smMapZ. (E) Dual color analysis of the movement of FtsZ and smMapZ. (F) Distribution of logD of single molecule tracking of smMapZ. D is the diffusion coefficient. The Gaussian fitting curve, the average D, and area of the two populations are shown. (G) Kymograph analysis of representative constrained molecules (left) and the localized relationship between the FtsZ ring in the newborn (upper) and the grown (lower) cells (right). (H) Snapshots of the time-lapse imaging of mMaple3_smMapZ. The smMapZ ring structure is marked by yellow arrows.

Single molecule tracking allowed us to measure the spatially specific dynamic features of targeted molecules. Here, we labeled Halotag fused smMapZ with low concentrations (62.5 pM) of HaloLigand-JF646 to monitor the motion of single smMapZ molecules. The results revealed two populations of smMapZ molecules: (i) stationary and (ii) diffusive (Figure 5F, Movie s10). In a few cases, we also observed one single smMapZ molecule switched between the two modes (Figure S5B). We analyzed the localization pattern of stationary smMapZ molecules in 62 cells. In 38 cells that exhibited newborn cell length or three FtsZ ring structures, the stationary smMapZ molecules co-localized with the FtsZ ring. In the rest 24 growing cells, the stationary smMapZ molecules localized at the cell equator of the future daughter cells (Figure 5G, Figure S5C). The low mobility of smMapZ molecules supports that smMapZ forms a stable ring structure. However, as most single molecules were bleached in 1-3 min, single molecule trajectories were insufficient to study the stability of the smMapZ ring over a longer period of time. Therefore, we labeled smMapZ with mMaple3, a photoconvertable fluorescent protein that converts from green fluorescence to red fluorescence under purple illumination.^29^ Using TIRF illumination, we photoconverted mMaple3-fused smMapZ molecules in the bottom portion of the ring using a 405 nm laser and monitored their movement every 3 min. The results showed that over a total duration of 21 min, neither the position nor the intensity of the photoconverted smMapZ molecules changed remarkably, indicating that the smMapZ ring is a stable structure and the exchange of smMapZ molecules occurs at a very slow rate (Figure 5H, Figure S5D). It is thus tempting to propose this stable smMapZ ring structure plays as a novel “track” role during cell division, which guides the dynamic treadmilling FtsZ filaments at the proper position to self-orgnize into Z-ring (Figure 6). To the extent of our knowledge, this is the first work that shows bacteria control the Z-ring formation by affecting the treadmilling of FtsZ filament.

**Figure 6.**
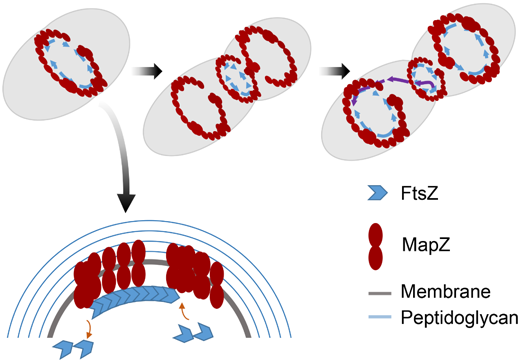
Model of smMapZ guiding the treadmilling of FtsZ. smMapZ forms a stable heterogeneous ring-like structure at the division site. The stable ring acts as a track for the treadmilling of FtsZ and, as the cell grows, this stable smMapZ ring moves to the future division site of daughter cells before the FtsZ ring. Newly formed FtsZ will move along the stable ring, and the translocated FtsZ filaments will participate in the new ring formation.

## Conclusion

A transmembrane protein MapZ-mediated FtsZ regulatory system was uncovered in *S. pneumoniae,* which was shown to be conserved in *Streptococcaceae.^11^* However, understanding the mechanism by which this system works is limited by the smattering of how MapZ regulates the FtsZ ring position. Here, our data suggest that a MapZ homolog in *S. mutans* forms a stable heterogeneous ring-like nanostructure at the division site, which may act as a track for the treadmilling of FtsZ filaments (Figure 6). Nevertheless, in spite of its essential role in guiding the position as well as the angle of the FtsZ ring, smMapZ is actually not required for FtsZ to form a ring. Our data reveal that the change of plane angle of the FtsZ ring often happened at the early stage of its formation but not in the constricting stage. Thus, we propose that the track role of smMapZ only functions at the early stage of FtsZ ring formation, and other proteins also participate in the stabilization of FtsZ treadmilling after Z ring formation, possibly including ErZA, a spectrin-like protein,^30^ which has been reported to interact with FtsZ and is required for correct localization of the FtsZ ring.^31^ Considering the similar functions of smMapZ with that of spMapZ, we believe that the “track” model of smMapZ does not only exist in *S. mutans,* but also in other MapZ-conserved bacteria.

Previously, two studies focused on the question that what regulates MapZ at division site, Sylvie Manuse *et al.* revealed the two sub-domains structure of the extracellular domain, indicating the two sub-domains could permit structure rearrangement to regulate MapZ binding to peptidoglycan.^15^ Renske van Raaphorst *et al.* claimed that the origin of chromosome replication (oriC), instead of MapZ, is crucial for division site selection.^16^ MapZ moves immediately following the position of oriC and is a ruler for the establishment of correct Z-ring plane angle.

Complementary to these studies, our study sheds light on how MapZ regulate the FtsZ ring position and provides new insights for the in-depth understanding of the MapZ-mediated FtsZ regulatory system. However, in addition to the “track” role, the MapZ ring may also serve as a “road fence” to restrict and guide the FtsZ treadmilling. In order to distinguish these two possibilities, the better understanding of the interaction between MapZ and FtsZ is required in future.

## Material and Methods

### Bacterial strains, plasmid and mutation construction, and culture conditions

*S*. *mutans* standard strain UA159 and all derived strains used in this study are listed in Table 1. To generate the *S*. *mutans* mutations (gene deletion and fluorescent protein fusion), we used a counter-selectable system called IFDC2 (provided by Zhoujie Xie). The transformation procedure was performed as previously described.^32^ Briefly, the overnight culture was grown in Todd-Hewitt medium supplemented with 0.3% yeast extract (THY; QinDao Wei si, China) was diluted (1:20) in the same medium and incubated for 2-3 h (OD600 0.2-0.3). Overlapping PCR reaction (5 μl) and 0.5 μl of CSP (1 μg/μl stock) were added to 500 μl of culture and incubated for 2 h. To select antibiotic-resistant colonies, brain heart infusion (BHI; Difco Laboratories, Detroit, MI, USA) agar plates were supplemented with 12-μg/ml erythromycin (Sigma-Aldrich, St. Louis, MO, USA). Selection of the second transformation was performed on BHI plates supplemented with 4mg/ml p-cl-Phe (Sigma-Aldrich). For snapshot imaging, *S*. *mutans* strains were aerobically cultured in THY at 37°C. For time-lapse imaging, the strains were cultured in a chemically defined medium^33^ supplemented with 0.3% yeast extract (C + Y). A final concentration of 10 mg/ml xylose was added to the culture medium to induce the expression of Pzx9 and Pzx10^34^ (provided by Jiezhou Xie).

### Sample preparation for snapshot imaging

An overnight culture grown in THY medium was diluted (1:100) in the same medium, and incubated at 37°C aerobically to OD600 0.4–0.5 (7 h). The culture was diluted again (1:100) in THY and grown at 37°C aerobically to OD600 0.1-0.2 (5 h). Cells were harvested by centrifugation at 1,500 × g for 3 min, washed three times with 1× phosphate-buffered saline (PBS), and resuspended in 1xPBS. The bacteria were loaded between a gel pad (1x PBS supplemented with 1.5% low melting agarose) and coverslip (Fisher 24 × 50, No. 1). For FtsZ imaging, spectinomycin was added to all cultures at a final concentration of 1 mg/ml. D (+) xylose (Sigma-Aldrich) was added before the last round of dilution at 10 mg/ml.

### Sample preparation for 3D-SIM imaging

JF_549_ dye (500 nM; provided by Luke Lavis) was added into the culture at OD600 0.1–0.2, incubated for 15 min at 37°C, washed once with 1 × PBS, fixed in 4% paraformaldehyde for 15 min at room temperature, and washed twice with 1 × PBS. Resuspended cells were loaded between the coverslip (Fisher 24 × 50, No. 1.5) and gel pad.

### Sample preparation for time-lapse imaging

#### EGFPsmMapZ and mMaple3_smMapZ

An overnight culture grown in THY medium was diluted (1:100) in C + Y medium. After two dilutions, the culture at OD_600_ 0.1-0.2 was added to a gel pad (C + Y medium supplemented with 1.5% low melting agarose) on a glass slide, and a coverslip was added. Rectangular 3M double-sided tape was used to seal the coverslip and glass slide.

#### HaloJF_646__smMapZ

JF_646_ dye (62.5 pM) was added into the culture at OD_600_ 0.1-0.2, incubated for 15 min at 37°C and washed once with pre-warmed fresh C + Y medium. Resuspended cells were loaded between the coverslip and the gel pad.

#### Imaging of FtsZ_mNeongreen

Spectinomycin was added to all cultures at a final concentration of 1 mg/ml. D (+) xylose was added at 10 mg/ml before the last round of dilution.

### Slide preparation

Coverslips were cleaned with Piranha solution (30% H2O2: 98% H2SO4 v:v = 1:3 at 90°C for 30 min), washed with Milli-Q water, stored in filtered Milli-Q water, and dried with high purity nitrogen before use Snapshot imaging of smMapZ and FtsZ

The samples were visualized on an N-STORM (Nikon, Tokyo, Japan) system equipped with a 100* oil TIRF objective (Nikon PLAN APO, 1.49 NA), Andor-897 EMCCD (Andor, Belfast, Northern Ireland), the perfect focus system (Nikon), a laser source (405, 488, 561, and 647 nm) and 1.5* magnification optics. Images were collected with Nis-Elements AR software (Nikon). For EGFP_smMapZ and mCherry_smMapZ, 300 frames were collected with a 100 ms exposure time. For FtsZ_EGFP, 50 frames were collected with a 100 ms exposure time.

#### Time-lapse imaging of EGFP_smMapZ and mMaple3_smMapZ

Time-lapse observation of EGFP_smMapZ was performed with a DeltaVision OMX SR (GE Healthcare, Little Chalfont, UK) system equipped with a 1.42 NA 60× oil objective (Olympus, Tokyo, Japan), laser source (405, 488, 561, and 642 nm), four sCOMS, and an environment control system. Using the OMX Acquisition software, the images were collected with a 60 ms exposure time, with 5 min intervals at 37°C.

Time-lapse imaging of mMaple3_smMapZ was performed with an N-STORM microscope equipped with a live cell instrument. A laser at 405 nm was used to photoconvert the mMaple3 to the red state with an initial exposure time of 300 ms. Then, a 561 nm laser was used to excite the red state mMaple3 with a 400 ms exposure time, and a 488 nm laser was used for the green state with a 300 ms exposure time.

#### 3D-SIM imaging of HaloJF549_smMapZ

Super-resolution 3D-SIM imaging was performed with a DeltaVision OMX SR system, with a 10 ms exposure time and 50% transmission. The 3D-SIM raw data were reconstructed with SoftWoRx 6.0 (Applied Precision, Issaquah, WA, USA) using a Wiener filter setting of 0.001 and a channel that specifically measured optical transfer functions. The immersion oil was optimized to 1.518.

#### Imaging of the dynamics of FtsZ_mNeongreen, mNeongreen_smMapZ and HaloJF646_smMapZ

The images were collected by near-TIRF illumination on an N-STORM microscope. Exposure time was 0.165 s, interval time was 0.495 s, and mNeongreen was excited by the 488 nm laser. The images were captured at room temperature. For the single molecular track of HaloJF646-smMapZ, exposure time was 0.1 s, and the interval time was 0.5 s. The 647 nm laser was used to excite the JF646 dye.

## Data analysis

### Localization of smMapZ

The averaged image stacks were obtained, and Snapshot analysis accomplished, with ImageJ software (NIH, Bethesda, MD, USA). Cell outlines and corresponding fluorescence signals were manually detected and analyzed using MicrobeTracker^35^ and custom MATLAB (MathWorks, Natick, MA, USA) codes. Information regarding the cell length of WT and ΔsmmapZ was extracted and the graph was generated in Microsoft Excel software (Microsoft Corp., Redmond, WA, USA). To compare the distance between the two outer rings, and between the smMapZ ring and the cell pole, we manually eliminated the cell in which only one ring structure could be observed, then the data were calculated with a custom MATLAB script and plotted in Excel. To sort the fluorescence intensity of the smMapZ truncations as a function of cell length, averaged images were deconvoluted using Huygens and subsequently analyzed using Coli-Inspector running under the ImageJ plugin ObjectJ (http://simon.bio.uva.nl/objectj/), as previously described.^15^

The analysis of periodic intensity fluctuations of the Z ring and treadmilling was performed in Fiji (http://fiji.sc/). First, drift correction of the time-lapse imaging was performed using series registration based on the Fiji plugin Descriptor. We selected a medium detection brightness and six-pixel for detection size; the type of detection was set as minima and maxima, and the rest were set as default values. After drift correction, a 3 × 3 pixel (~320 nm) region of interest was chosen inside the Z ring, and the Z-axis profile was plotted with Fiji to show the periodic intensity fluctuations. For analysis of FtsZ treadmilling, the images were resized to 26 nm/pixel with the bicubic method, and a segmented line (line width is 11) was drawn inside the Z ring, parallel to its long axis, and a multiple kymograph plugin was used to demonstrate movement. The velocity of treadmilling was calculated based on the kymograph using the macro “read velocities from tsp” as described in the manual.

### Single particle tracking

The Fiji plugin TrackMate (http://imagej.net/TrackMate) was used to analyze the trajectory of single molecules of smMapZ. We selected the LoG detector and set 0.4 μm as the estimated blob diameter. The LAP Tracker function, with a 0. 5 μm maximum distance for frame-to-frame linking, was employed to complete the tracking. The trajectory data were exported; further analysis of logD was carried out in R studio (R Development Core Team, Vienna, Austria) with a custom script. The two-status analysis of single trajectory was performed in MATLAB with the HMM-Bayes procedure,^36^ as previously described.

### smMapZ sequence analysis

To identify smMapZ, we selected spMapZ as the query sequence and used BLAST (http://blast.ncbi.nlm.nih.gov/Blast.cgi) to search the genome of *S*. *mutans*. The two sequences were aligned with SnapGene software (GSL Biotech LLC, Chicago, IL, USA). The transmembrane domain was predicted using TMHMM Server v. 2.0 (http://www.cbs.dtu.dk/services/TMHMM/). Network Protein Sequence Analysis^37^ (http://npsa-pbil.ibcp.fr) was used to predict the secondary structure of the cytoplasmic domain of smMapZ as previously described. In total, 150 protein sequences were selected to determine amino acid conservation using the Consurf web server^38^ (http://consurf.tau.ac.il/2016/). Prediction of sequence disordered tendency was performed with IUPred^39^ (http://iupred.enzim.hu/).

## Supporting information

Experimental data and strain list (PDF)

Movie s1. Directional movement of FtsZ inside the Z ring in WT cells (avi)

Movie s2. Directional movement of FtsZ inside the Z ring in *ΔsmmapZ* cells (avi) Movie s3. Translocation of FtsZ in WT cells (avi)

Movie s4. Translocation of FtsZ in *ΔsmmapZ* cells (avi)

Movie s5. Unconstrained movement of FtsZ in *ΔsmmapZ* cells (avi)

Movie s6. Unconstrained movement of FtsZ in *ΔsmmapZ* cells (avi)

Movie s7. The lifetime of constricting FtsZ ring in *ΔsmmapZ* cells (avi)

Movie s8. The lifetime of constricting FtsZ ring in WT cells (avi)

Movie s9. Timelapse of mNeongreen_smMapZ in WT cells (avi)

Movie s10. Single molecule tracking of smMapZ (avi)

Movie s11. The ring structure of HalojF_549__smMapZ (avi)

Figure S3. **Evolutionary tree analysis of smMapZ and spMapZ.** 151 MapZ homolog protein sequences in *Streptococcus* were selected by UniProt BLAST and NCBI protein blastp. Establish the evolutionary tree via Mrbayes (aamodelpr=mixed, 4,000,000 generations)

## ACKNOWLEDGEMENTS

This work was funded by the National Natural Science Foundation of China (No. 30973317, 31271423, and 21390412) and Peking University’s 985 Grant. We thank Zhoujie Xie (Institute of Microbiology, CAS) for his assistance in the construction of the mutations and Chunyan Shan (The Core Imaging Facility of Peking University) for her help on DeltaVision OMX. We thank He Yu, Professor Can Xie and ShuJin Luo (School of Life Sciences, Peking University) for their help with analysis of evolution tree.

## REFERENCES

1. Lowe, J.; Amos, L. A., Crystal structure of the bacterial cell-division protein FtsZ. Nature 1998, 391 (6663), 203–6.

2. Bi, E. F.; Lutkenhaus, J., FtsZ ring structure associated with division in Escherichia coli. Nature 1991, 354 (6349), 161–4.

3. Adams, D. W.; Errington, J., Bacterial cell division: assembly, maintenance and disassembly of the Z ring. Nature reviews. Microbiology 2009, 7 (9), 642–53.

4. Rowlett, V. W.; Margolin, W., The bacterial Min system. Current biology: CB 2013, 23 (13), R553–6.

5. Wu, L. J.; Errington, J., Nucleoid occlusion and bacterial cell division. Nature reviews. Microbiology 2011, 10 (1), 8–12.

6. Cho, H.; McManus, H. R.; Dove, S. L.; Bernhardt, T. G., Nucleoid occlusion factor SlmA is a DNA-activated FtsZ polymerization antagonist. Proc Natl Acad Sci US A 2011, 108 (9), 3773–8.

7. Hu, Z.; Mukherjee, A.; Pichoff, S.; Lutkenhaus, J., The MinC component of the division site selection system in Escherichia coli interacts with FtsZ to prevent polymerization. Proc Natl Acad Sci U S A 1999, 96 (26), 14819–24.

8. Thanbichler, M.; Shapiro, L., MipZ, a spatial regulator coordinating chromosome segregation with cell division in Caulobacter. Cell 2006, 126 (1), 147–62.

9. Treuner-Lange, A.; Aguiluz, K.; van der Does, C.; Gomez-Santos, N.; Harms, A.; Schumacher, D.; Lenz, P.; Hoppert, M.; Kahnt, J.; Munoz-Dorado, J.; Sogaard-Andersen, L., PomZ, a ParA-like protein, regulates Z-ring formation and cell division in Myxococcus xanthus. Mol Microbiol 2013, 87 (2), 235–53.

10. Willemse, J.; Borst, J. W.; de Waal, E.; Bisseling, T.; van Wezel, G. P., Positive control of cell division: FtsZ is recruited by SsgB during sporulation of Streptomyces. Genes & development 2011, 25 (1), 89–99.

11. Fleurie, A.; Lesterlin, C.; Manuse, S.; Zhao, C.; Cluzel, C.; Lavergne, J. P.; Franz-Wachtel, M.; Macek, B.; Combet, C.; Kuru, E.; VanNieuwenhze, M. S.; Brun, Y. V.; Sherratt, D.; Grangeasse, C., MapZ marks the division sites and positions FtsZ rings in Streptococcus pneumoniae. Nature 2014, 516 (7530), 259–262.

12. Holeckova, N.; Doubravova, L.; Massidda, O.; Molle, V.; Buriankova, K.; Benada, O.; Kofronova, O.; Ulrych, A.; Branny, P., LocZ is a new cell division protein involved in proper septum placement in Streptococcus pneumoniae. mBio 2014, 6 (1), e01700–14.

13. Banu, L. D.; Conrads, G.; Rehrauer, H.; Hussain, H.; Allan, E.; van der Ploeg, J. R., The Streptococcus mutans serine/threonine kinase, PknB, regulates competence development, bacteriocin production, and cell wall metabolism. Infection and immunity 2010, 78 (5), 2209–20.

14. Hussain, H.; Branny, P.; Allan, E., A eukaryotic-type serine/threonine protein kinase is required for biofilm formation, genetic competence, and acid resistance in Streptococcus mutans. Journal of bacteriology 2006, 188 (4), 1628–32.

15. Manuse, S.; Jean, N. L.; Guinot, M.; Lavergne, J. P.; Laguri, C.; Bougault, C. M.; VanNieuwenhze, M. S.; Grangeasse, C.; Simorre, J. P., Structure-function analysis of the extracellular domain of the pneumococcal cell division site positioning protein MapZ. Nature communications 2016, 7, 12071.

16. van Raaphorst, R.; Kjos, M.; Veening, J. W., Chromosome segregation drives division site selection in Streptococcus pneumoniae. Proc Natl Acad Sci U S A 2017, 114 (29), E5959–E5968.

17. Loose, M.; Mitchison, T. J., The bacterial cell division proteins FtsA and FtsZ self-organize into dynamic cytoskeletal patterns. Nature cell biology 2014, 16 (1), 38–46.

18. Yang, X.; Lyu, Z.; Miguel, A.; McQuillen, R.; Huang, K. C.; Xiao, J., GTPase activity-coupled treadmilling of the bacterial tubulin FtsZ organizes septal cell wall synthesis. Science 2017, 355 (6326), 744–747.

19. Bisson-Filho, A. W.; Hsu, Y. P.; Squyres, G. R.; Kuru, E.; Wu, F.; Jukes, C.; Sun, Y.; Dekker, C.; Holden, S.; VanNieuwenhze, M. S.; Brun, Y. V.; Garner, E. C., Treadmilling by FtsZ filaments drives peptidoglycan synthesis and bacterial cell division. Science 2017, 355 (6326), 739–743.

20. Lemos, J. A.; Quivey, R. G., Jr.; Koo, H.; Abranches, J., Streptococcus mutans: a new Gram-positive paradigm? Microbiology 2013, 159 (Pt 3), 436–45.

21. Garcia, P. S.; Simorre, J. P.; Brochier-Armanet, C.; Grangeasse, C., Cell division of Streptococcus pneumoniae: think positive! Current opinion in microbiology 2016, 34, 18–23.

22. Guo, F.; Tao, H.; Buss, J.; Coltharp, C.; Hensel, Z.; Jie, X., In Vivo Structure of the E. coli FtsZ-ring Revealed by Photoactivated Localization Microscopy (PALM). PloS one 2010, 5 (9).

23. Strauss, M. P.; Liew, A. T.; Turnbull, L.; Whitchurch, C. B.; Monahan, L. G.; Harry, E. J., 3D-SIM super resolution microscopy reveals a bead-like arrangement for FtsZ and the division machinery: implications for triggering cytokinesis. PLoS Biol 2012, 10 (9), e1001389.

24. Buss, J.; Coltharp, C.; Huang, T.; Pohlmeyer, C.; Wang, S.-C.; Hatem, C.; Xiao, J., In vivo organization of the FtsZ-ring by ZapA and ZapB revealed by quantitative superresolution microscopy. Molecular Microbiology 2013, 89 (6), 1099–1120.

25. Rowlett, V. W.; Margolin, W., 3D-SIM super-resolution of FtsZ and its membrane tethers in Escherichia coli cells. Biophys J 2014, 107 (8), L17–20.

26. Jacq, M.; Adam, V.; Bourgeois, D.; Moriscot, C.; Di Guilmi, A. M.; Vernet, T.; Morlot, C., Remodeling of the Z-Ring Nanostructure during the Streptococcus pneumoniae Cell Cycle Revealed by Photoactivated Localization Microscopy. mBio 2015, 6 (4).

27. Grimm, J. B.; English, B. P.; Chen, J.; Slaughter, J. P.; Zhang, Z.; Revyakin, A.; Patel, R.; Macklin, J. J.; Normanno, D.; Singer, R. H.; Lionnet, T.; Lavis, L. D., A general method to improve fluorophores for live-cell and single-molecule microscopy. Nature methods 2015, 12 (3), 244–50, 3 p following 250.

28. Bindels, D. S.; Haarbosch, L.; van Weeren, L.; Postma, M.; Wiese, K. E.; Mastop, M.; Aumonier, S.; Gotthard, G.; Royant, A.; Hink, M. A.; Gadella, T. W., Jr., mScarlet: a bright monomeric red fluorescent protein for cellular imaging. Nature methods 2017, 14 (1), 53–56.

29. Wang, S.; Moffitt, J. R.; Dempsey, G. T.; Xie, X. S.; Zhuang, X., Characterization and development of photoactivatable fluorescent proteins for single-molecule-based superresolution imaging. Proc Natl Acad Sci U S A 2014, 111 (23), 8452–7.

30. Cleverley, R. M.; Barrett, J. R.; Basle, A.; Bui, N. K.; Hewitt, L.; Solovyova, A.; Xu, Z. Q.; Daniel, R. A.; Dixon, N. E.; Harry, E. J.; Oakley, A. J.; Vollmer, W.; Lewis, R. J., Structure and function of a spectrin-like regulator of bacterial cytokinesis. Nature communications 2014, 5, 5421.

31. Jorge, A. M.; Hoiczyk, E.; Gomes, J. P.; Pinho, M. G., EzrA contributes to the regulation of cell size in Staphylococcus aureus. PloS one 2011, 6 (11), e27542.

32. Xie, Z.; Okinaga, T.; Qi, F.; Zhang, Z.; Merritt, J., Cloning-independent and counterselectable markerless mutagenesis system in Streptococcus mutans. Applied and environmental microbiology 2011, 77 (22), 8025–33.

33. van de Rijn, I.; Kessler, R. E., Growth characteristics of group A streptococci in a new chemically defined medium. Infection and immunity 1980, 27 (2), 444–8.

34. Xie, Z.; Qi, F.; Merritt, J., Development of a tunable wide-range gene induction system useful for the study of streptococcal toxin-antitoxin systems. Applied and environmental microbiology 2013, 79 (20), 6375–84.

35. Sliusarenko, O.; Heinritz, J.; Emonet, T.; Jacobs-Wagner, C., High-throughput, subpixel precision analysis of bacterial morphogenesis and intracellular spatio-temporal dynamics. Mol Microbiol 2011, 80 (3), 612–27.

36. Monnier, N.; Barry, Z.; Park, H. Y.; Su, K. C.; Katz, Z.; English, B. P.; Dey, A.; Pan, K.; Cheeseman, I. M.; Singer, R. H.; Bathe, M., Inferring transient particle transport dynamics in live cells. Nature methods 2015, 12 (9), 838–40.

37. Combet, C.; Blanchet, C.; Geourjon, C.; Deleage, G., NPS@: network protein sequence analysis. Trends in biochemical sciences 2000, 25 (3), 147–50.

38. Landau, M.; Mayrose, I.; Rosenberg, Y.; Glaser, F.; Martz, E.; Pupko, T.; Ben-Tal, N., ConSurf2005: the projection of evolutionary conservation scores of residues on protein structures. Nucleic acids research 2005, 33 (Web Server issue), W299–302.

39. Dosztanyi, Z.; Csizmok, V.; Tompa, P.; Simon, I., IUPred: web server for the prediction of intrinsically unstructured regions of proteins based on estimated energy content. Bioinformatics 2005, 21 (16), 3433–4.

